# DNA reconciles morphology and colouration in the drunk blenny genus *Scartichthys* (Teleostei: Blenniidae) and provides insights into their evolutionary history

**DOI:** 10.1101/2021.02.22.432327

**Authors:** E. Delrieu–Trottin, H. Hartmann Salvo, P. Saenz Agudelo, M. F. Landaeta, A. Perez Matus

**Affiliations:** UMR 5554 ISEM (IRD, UM, CNRS, EPHE), Univ Montpellier, Place Eugene Bataillon, 34095 Montpellier cedex 5, France; Museum fr Naturkunde, Leibniz Institute for Evolution and Biodiversity Science, Invalidenstr. 43, 10115 Berlin, Germany; CEFE, Univ Montpellier, CNRS, EPHE-PSL University, IRD, Campus CNRS 1919 route de Mende, 34293 Montpellier cedex 5 France; Subtidal Ecology Laboratory, Estacin Costera de Investigaciones Marinas, Departamento de Ecologa, Facultad de Ciencias Biolgicas Pontificia Universidad Catlica de Chile, Santiago, Casilla 114-D, Santiago, Chile; Instituto de Ciencias Ambientales y Evolutivas (ICAEV), Universidad Austral de Chile, Valdivia, Chile; Laboratorio de Ictioplancton (LABITI), Instituto de Biologa, Facultad de Ciencias, Universidad de Valparaso, Avenida Gran Bretaa 1111, Playa Ancha, Valparaso, Chile; Centro de Observacin Marino para Estudios del Ambiente Costero (COSTA-R), Universidad de Valparaso, Chile; Millennium Nucleus for Ecology and Conservation of Temperate Mesophotic Reef Ecosystem (NUTME)

**Keywords:** biogeography, Chile, integrative taxonomy, kelp forests, molecular phylogeny, species delimitation

## Abstract

The blenniids of the genus *Scartichthys* are one of the most common fishes of Central and South American Pacific coastal reefs. This being said, Scartichthys spp. remain difficult to identify in the field, and identification is particularly challenging across the ca. 6000 km where three of the four currently accepted species are known to occur in sympatry. A reason for this is that the main taxonomic characters from traditional taxonomy are indeed elusive. Additionally, At the same time, species can display multiple colour patterns in the field, depending on their ontogenetic stage, habitat association, and reproductive behaviour. Overall, molecular characterization is warranted to help address these issues. Here, we have used a novel approach to revise the genus by combining colouration, morphological, and molecular data of representative specimens of the four currently valid species and seven described colour patterns. From this, we show that only three of the four species should be considered as valid; *Scartichthys gigas* (Steindachner, 1876), *S. variolatus* (Valenciennes, 1836), and *S. viridis* (Valenciennes, 1836); while S. crapulatus Williams 1990 should be synonymized with *S. viridis*. In the same way, our analyses show that one of the colour patterns attributed so far only to *S. gigas* is characteristic of the juvenile stages of *S. viridis*. Our time-calibrated phylogeny shows that this genus is relatively young and that the estimated time of divergence between *Scartichthys gigas* and *S. viridis* is around 1.71 Ma. In comparison, the Desventuradas and Juan Fernandez Islands endemic *S. variolatus* diverged about 1.95 Ma. Our results help to clarify the taxonomy of Scartichthys.

**SIGNIFICANCE STATEMENT:** The blenniids of the genus Scartichthys are one of the most common fishes of Central and South American Pacific coastal reefs. Here we provide an updated phylogeny of this genus, comparing for the first time morphological, coloration, and molecular data in combination to resolve a 30-year-old discord among ecologists and taxonomists and discuss the potential underlying evolutionary processes that led to their presentday distribution.

## 1 INTRODUCTION

Species are the core units of any analysis in ecological, biogeographical, conservation, or evolutionary studies. Until the development of modern molecular biology tools, taxonomists described and named species solely on the basis of morphological characters, being the only tool available until the development of modern molecular biology tools (Teletchea, 2010). The recent development of high-throughput DNA sequencing technologies combined with global initiatives such as the Barcode of Life (Hebert & Gregory, 2005) have offered new tools and a framework to improve taxonomic descriptions and challenge traditional species descriptions. While the utility molecular approaches were once the subject of debated by taxonomists (Will et al., 2005), it is now recognized that the examination of different types of data (e.g. morphology, colouration, behaviour, molecules) can improve and accelerate the process of describing new species (Padial et al., 2010; Kekkonen & Hebert, 2014; Pante et al., 2015).

The blenniids of the genus *Scartichthys* are one of the most common fishes of the intertidal and shallow subtidal rocky environments of the Pacific coast of South America. Specifically, the distribution of these fishes stretches from western Panama to latitude 33S in Chile and also includes the Juan Fernndez Archipelago and the Desventuradas Islands (Stepien, 1990; Prez-Matus et al., 2017a, 2017b). Interestingly, Scartichthys are considered as blenniid giants as adults can be up to 300 mm in length. Additionally, they are known among local fishermen as borrachillas [drunk] because the consumption of the flesh of these fish leads to a sleepy or drunk feeling (Williams, 1990; Mndez-Abarca & Mundaca, 2016). Despite these interesting aspects of their ecology, Scartichthys blenniids remain difficult (Prez-Matus et al., 2007; Riquelme-Prez et al., 2019) or even impossible to identify at the species level in the field (Villegas et al., 2019). Identification is especially challenging across the 6000 km where three recognized species are known to occur in sympatry. Elusive characters such as the number of dental incisors (DI) and colour patterns described largely from preserved specimens (Williams 1990) are the main diagnostic characters used for the identification of the four currently valid species: Scartichthys variolatus (Valenciennes, 1836), *S. gigas* (Steindachner, 1876), *S. viridis* (Valenciennes, 1836), and S. crapulatus Williams 1990. Scartichthys variolatus is endemic to the islands of the Juan Fernndez Archipelago (33SL) and to the San Ambrosio and San Flix Islands (26SL). It is also the only Scartichthys species found on these islands. *Scartichthys gigas* (Steindachner, 1876) is distributed from Panama (9 NL) to northern Chile (Antofagasta, 23SL). In contrast to the three other species, *S. gigas* displays less than 73 DI and typically has 17 dorsal rays. *Scartichthys viridis* (Valenciennes, 1836) is geographically distributed from Peru (Independence Bay, 14SL) to Central Chile (Valparaso, 33 SL). Additionally, it displays more than 73 DI while the distinct colour patterns of preserved specimen (Tiny, dark-brown (rarely pale) spots on posterior half of body Williams (1990)) have commonly been used to describe it. Scartichthys crapulatus Williams 1990, is endemic to Central Chile and has only been reported from Central Chile (Barquito (26SL) and Valparaiso (33SL)) where it occurs in sympatry with *S. gigas* and *S. viridis*.

Using colouration patterns solely to identify species can be problematic and can lead to confusion in the field. A total of seven different colour patterns have been described for the four species of this genus based on preserved and live specimens. Mndez-Abarca & Mundaca (2016) described four different colour patterns for *Scartichthys gigas*; (1) the two-bar front head covered colour pattern, (2) the two-bar front head uncovered colour pattern, and (3) the reticulated bar-stained colour pattern, that all can be related to the reticulated colour pattern reported by Williams (1990) from preserved specimens of *S. gigas*. In addition to these three colour patterns, Mndez-Abarca & Mundaca (2016) described a new colour pattern for juveniles of *S. gigas*; (4) the uniform orange-brown colour pattern. This latter is problematic because it is very similar to one of the live colours patterns described by Williams (1990) for S. crapulatus, (5) the reddish-brown to golden colour pattern with orange-brown dots on the posterior half of the body. However, these orange-brown dots on the posterior half of the body are difficult to see in the field. (6) The dark-light bluish green colour pattern has been attributed to juveniles and adults of *S. viridis* (Mndez-Abarca & Mundaca, 2016). Finally, *S. variolatus* presents (7) a circular red spots in head and body colour pattern. The diversity of live colour patterns has been attributed to ontogeny, habitat association, and/or reproductive behaviour. Unfortunately, no explicit references have been made to classic morphological characters such as the number of DI for the new colour patterns described. It is worth noting that assessing these characters are particularly challenging for juveniles as teeth are still in developing during early ontogeny.

Finally, the recent phylogeny of blenniids (Hundt & Simons, 2018) based on five nuclear markers and which includes a representative of each of the four currently valid species is the first attempts to reconstruct the evolutionary history of Scartichthys (refer to Supplementary Figure S1 in Hundt & Simons (2018)). This molecular phylogeny confirms the monophyly of the genus Scartichthys. This molecular evidence and that of previous study (Hundt et al., 2014; Hundt & Simons, 2018) also confirm that Scartichthys is sister to Ophioblennius as formerly hypothesized by Williams (1990) using morphological data. Interestingly, Hundt & Simons (2018) retrieve minimal divergence between S. crapulatus and *S. viridis*, which parallels the concern expressed by Stepien (1990) regarding the validity of S. crapulatus.

The elusive diagnostic morphological characters of this group, the controversy regarding the validity of S. crapulatus, the lack of diagnostic molecular data, and the multitude of colour patterns described so far for the genus Scartichthys call for a reappraisal of the different diagnostic characters. This study aims at clarifying the taxonomy of Scartichthys using a novel approach including colouration, morphological, and molecular data in combination. Here, we present a reconstructed phylogeny including the different live colour patterns described so far for the four currently valid species of this genus to investigate the validity of the species described so far. We also reconstructed the evolutionary history of this genus and provide insights into the potential underlying evolutionary processes that led to the presentday distribution of Scartichthys species.

## 2 MATERIALS AND METHODS

### 2.1 Ethical statement sampling

Fishes were collected according to Chilean environmental laws using R.EX 2231, R.EX 556 and R.EX 1489 permits. Procedures for collection, maintenance, and analyses of fishes followed the international guidelines for animal experiments and using ethical permits from the Universidad de Valparaiso and Pontificia Universidad Catlica de Chile.

### 2.2 Taxon sampling

We analysed 66 specimens from eight locations, covering most of the geographic range of the four species that make up the Scartichthys species complex (Figure 1(a), Table S1).

**Figure 1:**
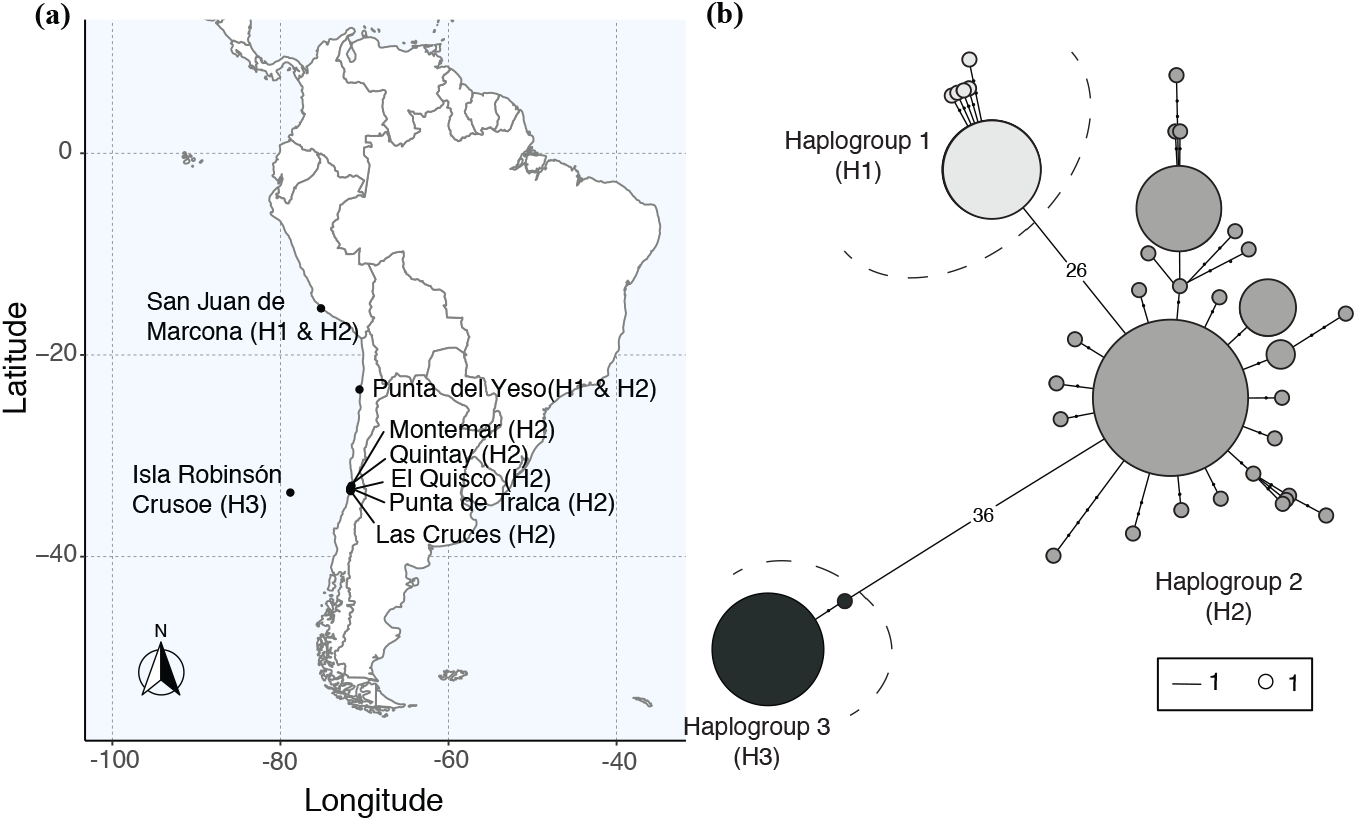
(a) Collection sites in Peru (San Juan de Marcona), continental Chile (Punta del Yeso, Montemar, Quintay, El Quisco, Punta de Tralca, Las Cruces), and Chilean oceanic islands (Isla Robinson Crusoe), H1, H2, and H2 referring to the haplogroup found in each site, and (b) Haplotype network for the genus Scartichthys; each circle corresponds to a unique sequence (i.e., haplotype) and size of circles indicates haplotype frequency.

For this study, 55 specimens were sampled between 0 and 15 m depth using hand nets from December 2018 to February 2019. Also, 11 specimens (early juveniles) were captured between Montemar, 33SL, and El Quisco, 33SL, between September 2015 and February 2017 using Ecocean light traps (CARE, Ecocean, Montpellier, France). All light trap samples were preserved in 96% EtOH (specimens from Daz-Astudillo et al. (2019). Following bioethical standards, all specimens were euthanized with an overdose of benzocaine before preservation. A small piece of pectoral fin tissue from each specimen was preserved in 96% EtOH at −20C.

### 2.3 Morphological analyses

Counts of incisor teeth and dorsal fin rays followed Williams (1990) and were taken using a Leica model EZ4 binocular. Following Williams (1990) and Mndez-Abarca & Mundaca (2016), specimens were photographed using digital cameras (Nikon D90 and Canon EOS T5) upon collection or in the laboratory to identify their fresh colouration patterns. The total counts of dentary incisors (DI) are diagnostic for only two of the four species of this genus (*S. gigas* and *S. variolatus*). Additionally, the seven live colour patterns in life currently reported so far allow the four currently described Scartichthys to be distinguished in the field. It should be noted that the morphological data (colouration) was collected for specimens collected in San Juan de Marcona (SJM), Per is incomplete. No morphological data could be obtained from specimens preserved in alcohol upon capture (Table S1).

### 2.4 Molecular analyses

#### 2.4.1 DNA Extraction, amplification and sequencing

Whole genomic DNA was extracted from fin tissue preserved in 96% EtOH. DNA extraction was performed following the HotSHOT method (Truett et al., 2000), using 50mM NaOH and 1M Tris-HCL. For each specimen, we amplified a 652 bp fragment of the mitochondrial gene coding for cytochrome C oxidase subunit I (COI). The primers F2 and R2 designed by Ward et al. (2005) were used. Fragments were amplified using PCR protocols as described by Williams et al. (2012), Modifications in the final reactions (10 l) which contained 5 l of KAPAG Fast Multiplex Mix multiple mixing solution (KAPA2G Fast HotStart DNA Polymerase (1 U per 25 L reaction), KAPA2G Buffer A (1.5X at 1X), dNTPs (0.2 mM each dNTP at 1X), MgCl2 (3 mM at 1X) and stabilizers), 1 l of a mixture of F2 and R2 primers (2mM each primer), 3.0 l H2O, and 1 l of genomic DNA. After PCRs were complete, 1 l of each PCR product (mixed with 1 l of Red Gel dye) was separated by electrophoresis on a 1% agarose gel at 100 V for 30 minutes and visualized with a UV transilluminator. When the PCR products showing a clear and unique band of the correct expected length, all PCR products were purified by adding 1.3 l of alkaline exonuclease phosphatase; purification was conducted in a thermal cycler at 37C for 60 min and then at 85C for 15 min. Subsequently, sequencing was performed bidirectionally with the same PCR primers using the BigDye Terminator v3.1 cycle sequencing kit and an ABI 3500 XL Applied Biosystems sequencer. Sequences were aligned with Clustal W (Thompson et al., 1994) and edited using GENEIOUS 9.0.5 (http://www.geneious.com, (Kearse et al., 2012)). All generated sequences were deposited in GenBank (Accession numbers: OL413590 - OL413655).

#### 2.4.2 Genetic distances and sequence-based species delimitation analysis

To visualize the relationships between haplotypes of Scartichthys spp. among the different sampling localities, we first constructed a haplotype network using the haplonet function of the package pegas (Paradis, 2010) in the R statistical environment (R Core Team, 2020). We then implemented Neighbour-Joining (NJ), using the software package MEGA 6 (Tamura et al., 2013). Confidence in topology was evaluated by bootstrap analysis with 1000 replicates (Felsenstein, 1985). Maximum Likelihood and Bayesian inference methods were used to perform sequence-based species delimitation analysis. We used sequences of Ophioblennius macclurei (KF930203), Cirripectes variolosus (MH707881), Cirripectes polyzona (HQ168554) and Exallias brevis (MF409572) downloaded from GenBank to root the trees in all analyses. As each sequence-based species delimitation method is susceptible to pitfalls, we used a 50% consensus among the five different methods we implemented as a robust delimitation scheme (Kekkonen & Hebert, 2014; Hubert & Hanner, 2015; Kekkonen et al., 2015): (1) Automatic Barcode Gap Discovery (ABGD; Puillandre et al., 2012) available at https://bioinfo.mnhn.fr/abi/public/abgd/, (2) single (sPTP) and (3) multiple rate version (mPTP)of the Poisson Tree Process (PTP, Zhang et al., 2013) available at https://mptp.h-its.org//tree and (4) single (sGMYC) and (5) multiple rate version (mGMYC) of General Mixed Yule-Coalescent (GMYC) as implemented in the R package Splits 1.0-19 (Fujisawa & Barraclough, 2013). The ABGD analysis requires a DNA alignment as the input; the PTP analyses require a ML tree as the inputs, and GMYC analyses require an ultrametric tree as the inputs. The Maximum Likelihood (ML) analysis was performed using the online version of IQTREE (Minh et al., 2013; Nguyen et al., 2015) available at http://iqtree.cibiv.univie.ac.at (Trifinopoulos et al., 2016). ModelFinder implemented in IQ TREE was used to assess the best model of evolution using the Bayesian Information Criterion (BIC) prior to the construction of the ML tree (Kalyaanamoorthy et al., 2017). The ultrafast bootstrap approximation (UFboot) (Minh et al., 2013) and the SHlike approximate likelihood ratio test (SHaLRT), both with 1,000 bootstrap replicates (Guindon et al., 2010) were conducted to evaluate the reliability of the nodes. Finally, the ultrametric and fully resolved tree needed to conduct the GMYC analyses was reconstructed using the Bayesian approach implemented in BEAST 2.5.2. Duplicated sequences were pruned prior to reconstructing the ultrametric tree. Two Markov chains of 10 million generations were run independently using a Yule pure birth model tree prior, a strict-clock model of 1.2% of genetic distance per million years. Trees were sampled every 1,000 states after an initial burn-in period of 1 million generation. Both runs were combined using LogCombiner 2.5.2, and the maximum credibility tree was constructed using TreeAnnotator 2.5.2 (Bouckaert et al., 2019).

### 2.5 Multilocus time-calibrated phylogeny

A time-calibrated phylogeny (BI) of *Scartichthys* spp. was constructed using the software BEAST2 2.5.2 (Bouckaert et al., 2019). Six molecular markers for each species were included; specifically one mitochondrial and five nuclear regions were used. For the mitochondrial marker, we used *S. viridis, S. gigas* and *S. variolatus* representative COI sequences produced in this study. In contrast, for these three species we used nuclear sequences (ENC1, myh6, sreb2 and tbr1) produced by Hundt & Simons (2018) (Table S2). We did not include Scartichthys crapulatus in this analysis as all our previous results indicate that S. crapulatus and *S. viridis* actually represent the same species (see Results section). Blenniids are relatively rare in the fossil record (Bannikov 1998), but see Liu et al. (2018) for a recent review, preventing their use as a reliable calibration point. Deep secondary calibrations have thus been generally used for blenniids (e.g. Lin & Hastings, 2013; Liu et al., 2018), which can lead to overestimation of divergence times among taxa that have recently diverged (Ho et al., 2008) or in such small scale survey. We thus chose to use an informative prior for the evolutionary rate of COI of 1.2 x10-8 substitution per site per years commonly used for fishes for this marker (e.g Bermingham, McCafferty, & Martin, 1997; Lessios, 2008;Tea et al.,2019). We assumed a strict clock for each of the six markers, with the relative rates of ENC1, myh6, ptr, sreb2 and tbr1 being inferred in our analyses. Additionally, a BirthDeath model with a chain length of 30 million generations was used as the tree prior. ModelFinder implemented in IQ TREE was used to assess the best model of evolution for each marker using the Bayesian Information Criterion (BIC). Trees and parameters were sampled every 3000 generations, and the first 10% of the samples were discarded as burn-in. We assessed the convergence and appropriate burnin of each analysis using TRACER 1.5 (Drummond & Rambaut, 2007). Three independent analyses were run to ensure convergence. A maximum clade credibility tree was constructed using TreeAnnotator 2.5.2 (Bouckaert et al., 2019) to get median ages and 95% highest posterior density (HPD) intervals for each node. The 95% HPD represents the smallest interval that contains 95% of the posterior probability and can be loosely thought of as a Bayesian analog to a confidence interval (Gelman et al., 2013).

## 3 RESULTS

The total length (TL) of specimens collected ranged from 50 to 246 mm (Figure 2, Table S1).

**Figure 2:**
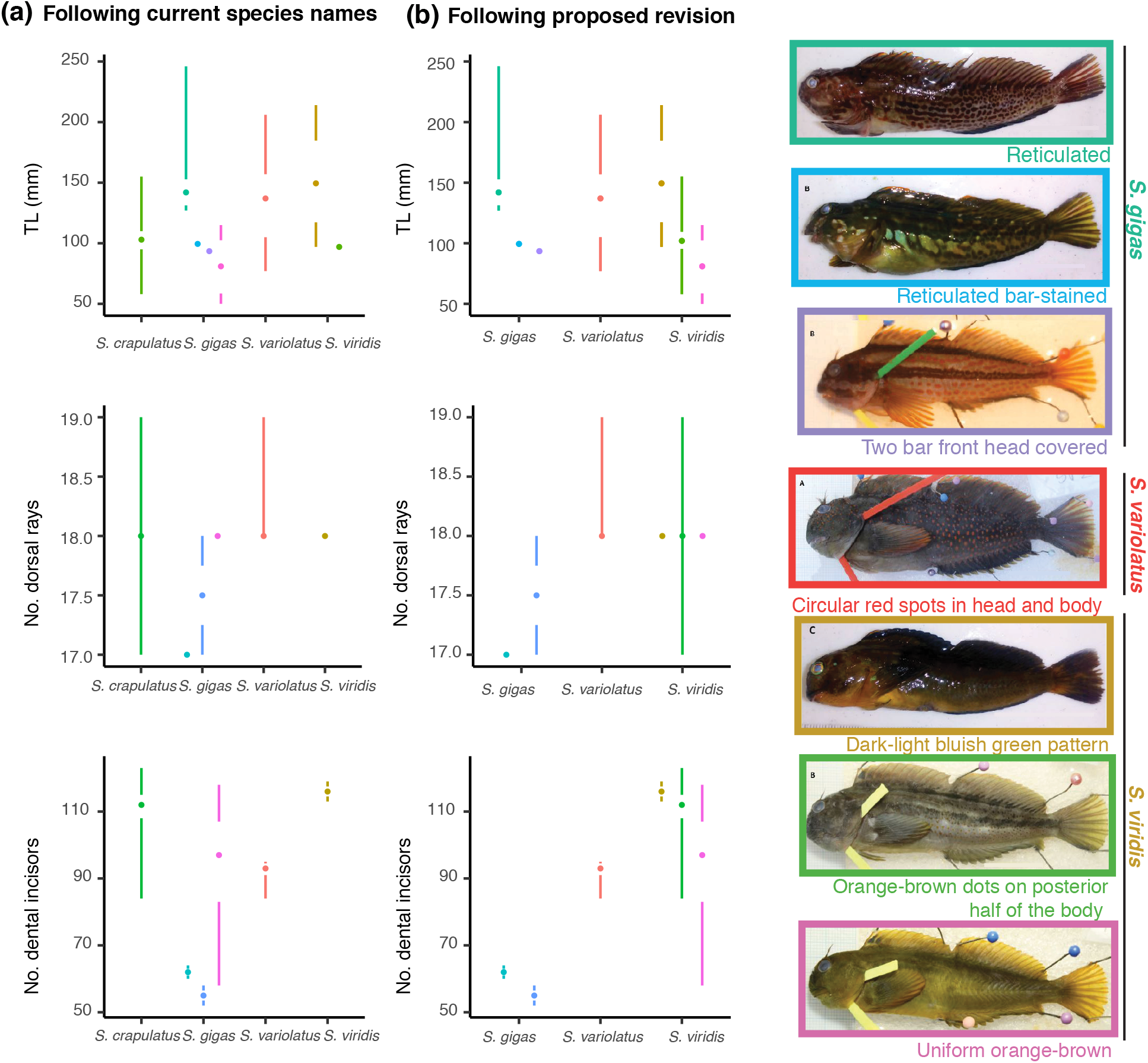
Tufte boxplot representing Total length (TL (mm)) (top), number of dorsal rays (middle) and number of dental incisors (bottom) for each colouration pattern when classified (a) following current species names, and (b) following proposed revision for the seven colour patterns observed for the genus Scartichthys. In each panel and for each species, the median is represented by a point, the interquartile range (25-75%) by a gap between vertical lines and the lines indicate the range of values below and above quartiles.

Specimens displaying the uniform orange-brown colour pattern and the orange-brown dots on posterior half of the body colour pattern were the smallest, with mean size 80 mm ( 24 mm) and 106 mm ( 20 mm), respectively. In contrast, specimens displaying the reticulated patterns were the largest with a mean size 156 mm ( 44 mm). The number of dorsal rays retrieved for the reticulated bar stained colour pattern and the two bar front head covered colour pattern were typically 17 while the five other colour patterns displayed typically 18 (Figure 2). The number of dentary incisors (DI) was counted for 66 of the specimens and ranged from 52 to 123 (Figure 2). Among the three-colour patterns attributed to *Scartichthys gigas*, only the reticulated colour pattern and the two-bar front head covered colour pattern displayed a number of DI in accordance with the diagnostic description. Specimens with the uniform orange-brown colour pattern presented up to 118 DI. Interestingly, red/orange spots (dots) that are usually a diagnostic character of S. crapulatus were found on specimens with the uniform orange-brown colour pattern and all specimens of dark-light bluish-green colouration when observed under the binocular (see Figure S1). Scartichthys variolatus displayed a unique colour pattern (circular red spots on head and body) and DI number (80-93). Both characters can be used to distinguish this island species from the other continental species. We retrieved DI numbers in accordance with the diagnostic report. Finally, the colour pattern is the only character distinguishing S. crapulatus from *S. viridis*; we retrieved similar DI numbers for both colour patterns described so far for these two species (Figure 2). It is worth noting that the range of values retrieved for specimens displaying the uniform orange-brown colour pattern attributed so far only to *S. gigas* correspond with those of S. crapulatus and *S. viridis*.

Specimens displaying the uniform orange-brown colour pattern and the orange-brown dots on posterior half of the body colour pattern were the smallest, with a mean size 80 mm ( 24 mm) and 106 mm (20 mm), respectively. In contrast, specimens displaying the reticulated patterns were the largest with a mean size 156 mm ( 44 mm). The number of dorsal rays retrieved for the reticulated bar stained colour pattern and the two bar front head covered colour pattern were typically 17 while the five other colour patterns displayed typically 18 (Figure 2). Among the three colour patterns attributed to *Scartichthys gigas*, only the reticulated colour pattern and the two-bar front head covered colour pattern displayed a number of DI in accordance with the diagnostic description. Specimens with the uniform orange-brown colour pattern presented up to 118 DI. Scartichthys variolatus displayed a unique colour pattern (circular red spots in head and body) and DI number (80-93). Both characters can be used to distinguish this island species from the remaining continental species. We retrieved DI numbers in accordance with the diagnostic report. Finally, the colour pattern is the only character distinguishing S. crapulatus from *S. viridis;*we retrieved similar DI numbers for both colour patterns described so far for these two species (Figure 2). It is worth noting that the range of values retrieved for specimens displaying the uniform orange-brown colour pattern attributed so far to *S. gigas* correspond with those of S. crapulatus and *S. viridis*.

For the molecular analyses, we first worked from an alignment of 652 base pairs from the mitochondrial COI region. While specimens of all four currently four valid Scartichthys species were sampled, our molecular analysis only shows the existence of three well-supported and highly divergent clades. The haplotype network analysis shows 3 distinct groups, separated by 36 and 26 mutations, respectively: tthe first group is composed of all Robinson Crusoe specimens, the second group is composed of specimens collected in the two northernmost sampling localities (San Juan de Marcona (Peru) and Antofagasta (Chile)), and finally the third group is composed of specimens caught in continental Chile (Figure 1(b)). The best nucleotide substitution model using the Bayesian information criterion (BIC) was HKY+I+G. The NJ and ML approaches produced the same tree topologies with strong bootstrap support (Figure 3). Similarly, the Bayesian analysis produced the same tree topology (Figure 3) across all three runs with high posterior probabilities (PP) and parameters that reached effective sample sizes higher than 200.

**Figure 3:**
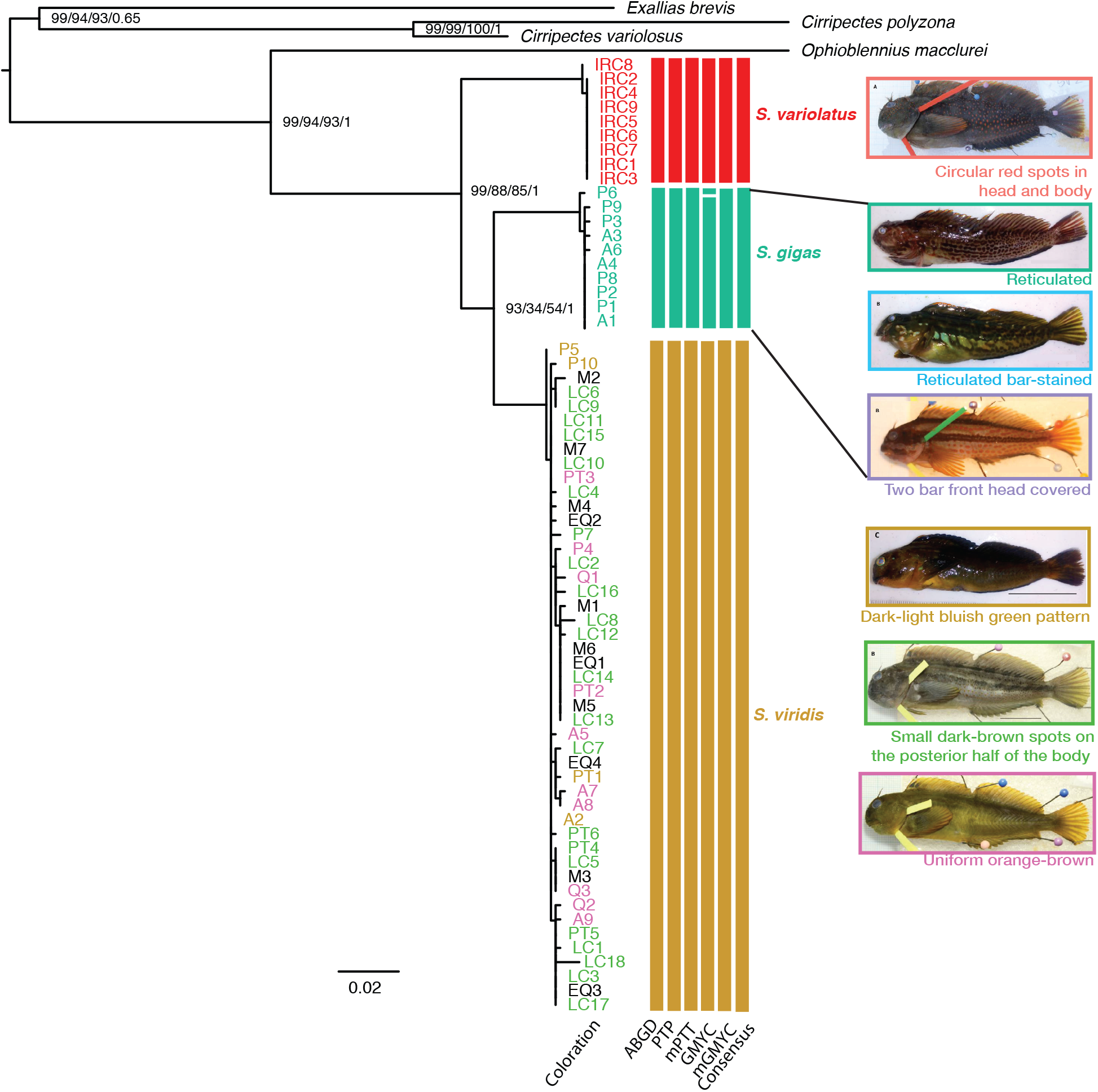
Maximum Likelihood tree for the 66 COI sequences generated in this study. Numbers on nodes denote bootstrap values with 1000 replicates (NJ analysis), UFboot values with 1000 replicates and the SH-aLRT also with 1000 bootstrap replicates (ML analysis); and Posterior Probabilities (BI). Vertical bars on the right side of the ML tree correspond to DNA-based species delimitation schemes derived from 5 methods (with corresponding abbreviations for each method at the bottom of each bar) and the resulting consensus delimitation scheme. Colouration code of tip labels denotes the different identifications based on morphology / colouration; no identification was possible for specimens in black (post larvae, see Table S1).

All three methods revealed that Scartichthys is a monophyletic group composed of three well supported and highly divergent clades: (1) The first clade is composed of all specimens from Robinson Crusoe Island identified as *S. variolatus*. (2) The second clade is composed of 11 out of the 20 specimens identified as *S. gigas*, representing all specimens with the reticulated and the two-bar front head covered colour pattern. (3) The third clade is composed of not only all specimens identified as *S. viridis* but also of all specimens identified as S. crapulatus and all specimens displaying the uniform orange-brown colour pattern described so far as a juvenile colour pattern for *S. gigas* (Figure 3). Species delimitation analyses provided a concordant number of Molecular Operational Taxonomic Units (MOTUs) among the different methods: three for ABGD, PTP, mPTP, GMYC and four for mGMYC. Thus, a a consensus delimitation scheme of three MOTUs was used thus forward. It is worth noting that the fourth MOTU delimited by mGMYC corresponds to a single specimen among the *S. gigas* clade presenting a reticulated bar-stained colour pattern.

The same topology was retrieved in our timecalibrated phylogeny based on six markers, one mitochondrial (COI) and five nuclear (ENC1, myh6, ptr, sreb2 and tbr1) (Figure 4).

**Figure 4:**
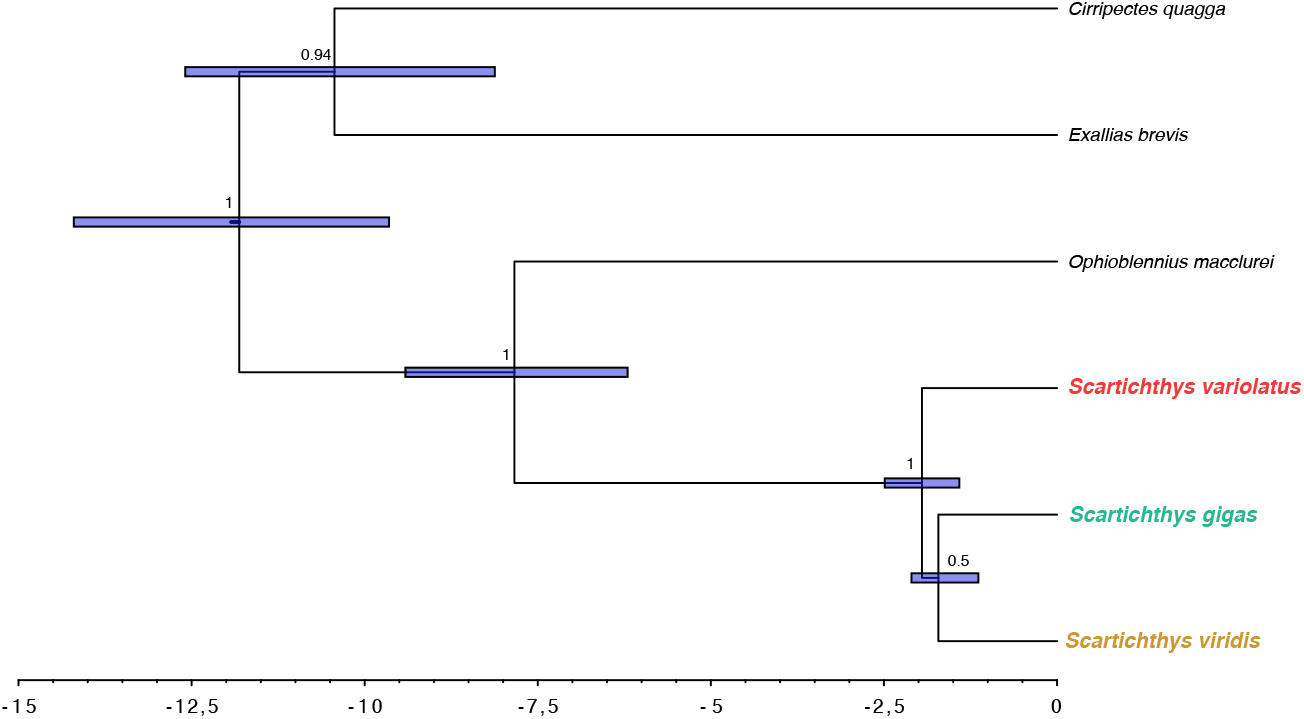
Time calibrated phylogeny for the genus Scartichthys. The Bayesian Maximum Clade Credibility Tree is based on 6 partitions (1 mitochondrial gene and 5 nuclear genes) and includes posterior probabilities (PP) and confidence intervals of divergence time estimates (95% HPD). Scale bar in Myr.

All nodes but one (*S. gigas S. viridis*; 0.50) were well supported (above 0.9). We found that Scartichthys and Ophioblennius diverged 7.84 Ma (6.20 9.41, 95% HPD). The estimated time of divergence between the *S. variolatus* (Clade 1) and the other Scartichthys spp. is approximately 1.95 Ma (1.41-2.49, 95% HPD; Figure 4) while *S. gigas* (Clade 2) and *S. viridis* (Clade 3) diverged 1.71 Ma (1.00 - 1.97, 95% HPD, Figure 4).

## 4 DISCUSSION

The present study represents an updated phylogeny of the Scartichthys genus. Using a combination of morphological, colouration, and molecular evidence, we show that this genus is composed of three species. Here, our integrative approach reveals that the colour pattern used to diagnose S. crapulatus and the uniform orange-brown colour patterns recently attributed to juveniles of *S. gigas* are both juvenile colour patterns of *S. viridis*. Finally, we show that the diversification of this genus is relatively recent beginning around 1.95 Ma.

A diversity of information has been produced so far to describe and characterize species of the genus Scartichthys. Here, for the first time, we have used this broad array of evidence with an integrative approach to clarify the taxonomy of Scartichthys. Five lines of evidence taken together indicate that the colour pattern small dark-brown spots on the posterior half of the body used to characterize Scartichthys crapulatus is also the colour pattern of juveniles of *S. viridis*. Additionally, the uniform orange-brown colour pattern is characteristic of *S. viridis* juveniles not of *S. gigas* as previously proposed. These conclusions are based on the following:

1. Size: The small dark-brown spots on the pos-terior half of the body and the uniform orange-brown specimens observed and collected in the field and collected measured respectively 106 mm ( 20 mm) and 80 mm ( 24 mm), respectively. This is substantially smaller than the typical size of what Scartichthys adults. These two colour patterns are thus likely characteristic of juveniles. Stepien (1990) first suggested that the colour pattern attributed to S. crapulatus (small dark-brown spots on the posterior half of the body) was actually one of the juvenile forms of an already existing species (*S. viridis*). Indeed, both juveniles and adults of *S. viridis* can be found displaying a dark-light bluish green pattern;
2. Colouration: We found small dark-brown spots on the posterior half of the body of not only adult specimens presenting the dark-light bluish green pattern characteristic of *Scartichthys viridis*, similarly to Mndez-Abarca & Mundaca (2016), but also on specimens displaying the uniform orange-brown pattern (so far attributed to *S. gigas* juveniles). The main diagnostic characteristic of S. crapulatus is thus not conclusive as three colour patterns share these spots. Interestingly, Williams (1990) mention that live colour pattern of S. crapulatus are highly variable, with Body colours [ranging] from green or reddish brown to golden with small brownish orange or brown spots on posterior half of body and on segmented-ray portion of dorsal fin. It is possible that specimens presenting an uniform orange-brown patterns were actually identified as S. crapulatus at that time. It should be emphasized that the two bar front head uncovered colour pattern described by Mndez-Abarca and Mundaca (2016) and attributed specifically to *S. gigas* corresponds to the Orange-brown dots on posterior half of body colour pattern described by Williams (1990) and is used in the present study as a character representative of *S. viridis* (Figure 5). While Mndez-Abarca & Mundaca (2016) did not mention orange-brown dots, we found that specimens with these two colour patterns have the two dark bars and the front head uncovered, and it is likely that dots were not easily noticeable on live specimens (see live specimens in Figure 5).
3. Dental incisors and dorsal rays: The three colour patterns sharing the small dark-brown spots on the posterior half of the body (dark-light bluish green, uniform orange-brown and orange-brown dots on posterior half of the body patterns) also share a similar number of DI. Interestingly, the number of DI of these fishes is clearly higher than the number of DI observed in Scartichthys gigas (Figure 2) and generally higher than the number of DI observed in *S. variolatus*. They also share a similarly higher number of dorsal rays, typically 18, compared to the number of dorsal rays typically observed for *S. gigas*. The more variable number of DI (109 8 but as low as 58 in small specimens) and dorsal rays (typically 18, but as low as 17) for the uniform orange-brown fishes retrieved here might have led Mndez-Abarca & Mundaca (2016) to attribute this colour pattern to *S. gigas*.
4. Molecular data: Molecular analyses showed a lack of reciprocal mtDNA monophyly for the small dark-brown spots on the posterior half of the body, the uniform orange-brown, and dark-light bluish green patterns. Indeed, phylogenetic analyses (distance-based, Maximum Likelihood, and Bayesian) and sequence-based delimitation methods all showed that these three-colour patterns were part of a single clade.
5. Distribution: Our extensive sampling allowed us to find the juvenile uniform orange-brown patterns in a geographic region where *Scartichthys gigas* has not yet been recorded in central Chile (Valparaiso region). No *S. gigas* adults have been observed excluding the possibility of a range extension of *S. gigas* and reinforcing the hypothesis that the uniform orange-brown is one of the juvenile colour patterns in *S. viridis*. The same reasoning applies for the two bar front head uncovered colour pattern. Specimens displaying the uniform orange-brown colour pattern are frequently observed in the subtidal kelp, Lessonia trabeculata, down to 20 m depth. Thus, we hypothesize that this colouration is involved in camouflage (Gaither et al., 2020). In line with Gaither et al. (2020), different colour morphs of *S. viridis* have been observed together inhabiting the same environments. This colour polymorphism could thus be related to either juvenile stages or microhabitat preferences as both colour morphs are associated with algae (Prez-Matus et al., 2017; Riquelme-Prez et al., 2019). Our study also allowed us to depict the evolutionary history of the Scartichthys genus. The phylogenetic analyses retrieved the monophyly of the genus Scartichthys, as previously proposed using a single representative per species (Hundt & Simons, 2018). Based on multiple specimens per colour patterns as well as currently valid species, we show here that the extremely shallow divergence observed by Hundt & Simons (2018) between *S. viridis* and S. crapulatus can be attributed to intraspecific divergence. Indeed, all of our analysis converged on the existence of three and not four well-supported clades within Scartichthys and resulted in trees presenting the same topology whether analyses were performed using COI only or using the mitochondrial marker and the five nuclear markers. The topology retrieved here differs from the one previously proposed based solely on nuclear markers. In that the previous study, Hundt & Simons (2018) have placed *Scartichthys gigas* as the earliest diverging species, while our results place Scartichthys variolatus in this position. Mitochondrial markers such as COI indeed often possess numerous informative sites to untangle relationships within species complexes but are more prone to saturation and homoplasy than conserved nuclear markers. These conserved markers are often preferred to untangle relationships between genera (Clabaut et al., 2005; Dornburg et al., 2014) and are poorly informative for the relatively young Scartichthys species complex. Scartichthys is sister to *Ophioblennius*,which are found in the Atlantic (*O.atlanticus*, *O. macclurei*, and *O. trinitatis*) and the Eastern Pacific (*O. clippertonensis* and *O. steindachneri*) (Froese & Pauly, 2019). Our estimates for the divergence between these two genera (7.84 Ma (6.20 9.41, 95% HPD)) are in agreement with the first estimations by Liu et al. (2018) (around 12 Ma (5 19 Ma) who used the crown Blenniidae (66 Mya) as their secondary calibration. *Scartichthys* and *Ophioblennius* genera diverging from the Indo-Pacific *Cirripectes* and *Exallia* via eastward is a likely dispersal hypothesis for the origin of *Scartichthys* in the eastern Pacific (Duncan et al., 2006; Hou & Li, 2018). Based on estimates of geographic range distributions, three latitudinal biogeographic regions have been described for the continental Chilean species that inhabit intertidal habitats. Interestingly, these also coincide with the ecoregions delineated within the Warm Temperate Southeastern Pacific Province (Spalding et al., 2007). These ecoregions include the central Chile that ranges from 20 to 36 SL, the Araucanian ecoregion from 39 to 43 SL, and the Chiloense region from 45 to 55 SL (Spalding et al., 2007; Navarrete et al., 2014). Here, the two closely related continental species *Scartichthys viridis* and *S. gigas* are both found within Central Chile. However *Scartichthys gigas* occurs from 9 N to 23 S, and *S. viridis* is found from 14 S to 33 S. The two species only co-occur from 14S to 23S. They differ indeed in their success to colonize warmer waters; *S. viridis* is restricted to the Humboldt Current system and found only up to 14 S. *Scartichthys gigas*, on the other hand, can be found not as south as *S. viridis* but as north as 9 N, outside of the Humboldt Current system, within the Tropical Eastern Pacific Province. The Desventuradas and Juan Fernandez Islands represent a distinct biogeographic unit (Dyer & Westneat, 2010) and a hotspot of endemism with more than 42 % of the species observed in their waters occurring nowhere else in the world (Dyer & Westneat, 2010). Their origin and the processes that led to their emergence remain somehow a mystery. Biogeographic analyses based on species distribution estimates have shown that the Desventuradas Islands can be grouped with Easter Island, and Sala y Gomez (Kulbicki et al., 2013). Interestingly, this cluster of islands is related to the Hawaiian Archipelago and the South Western Pacific Ocean province, which extends from western Australia all the way to the Kermadec islands (Kulbicki et al., 2013). Phylogenetic analyses including Juan Fernandez endemics are scarce and showed that Juan Fernandez endemic species are mostly related to either southwest Pacific species or southern Pacific species. More specifically, the Juan Fernandez endemics *Chironemus bicornis* (Steindachner, 1898), *C. delfini* (Porter, 1914), *Amphichaetodon melbae* Burgess & Caldwell, 1978 and *Girella albostriata* Steindachner, 1898 are closely related to Australian / New Zealand / Lord How Island species (Burridge et al., 2006; Cowman & Bellwood, 2011; Gaboriau et al., 2018; Delrieu-Trottin et al., 2019; Beldade et al., 2020) while *Pseudolabrus gayi* (Valenciennes, 1839) which occurs from Australia to Easter Island is closely related to the southern Pacific *P. fuentesi* (Regan, 1913) (Delrieu-Trottin et al., 2019). Ourphylogenetic analyses showed however that Juan Fernandez endemic species could also be related to continental Chilean species. The distinct origins found so far for these fishes call for a more extensive study including a larger number of Juan Fernandez endemic species. The time tree produced here shows that the divergence time of Scartichthys variolatus is around 1.95 Ma (1.41-2.49, 95% HPD). As such, this species can be considered as a Neoendemic, as it is younger than the geological age of the Juan Fernandez and Desventuradas Islands (Santa Clara: 5.8 Ma, Robinson Crusoe: 3.7 Ma (Clouard & Bonneville, 2005; Lara et al., 2018)). Since the Desventudas Islands are younger, with San Ambrosio being 2.9 million years old (Clouard & Bonneville, 2005), it is likely that speciation occurred on Juan Fernandez Island following a colonization of the Desventudas Islands. Similar divergence times have been retrieved for *Pseudolabrus gayi*, whose divergence dates around 2.35 Ma (1.34-3.63, 95% HPD) (Delrieu-Trottin et al., 2019). In contrast, the divergence of *Amphichaetodon melbae* (7.26 Ma (4.23-10.97, 95% HPD)) is likely older than the geological age of the Juan Fernandez islands (Delrieu-Trottin et al., 2019). A comprehensive survey of the divergence estimates of Juan Fernandez endemics would provide a better insight into the origin of this unique fauna.

**Figure 5:**
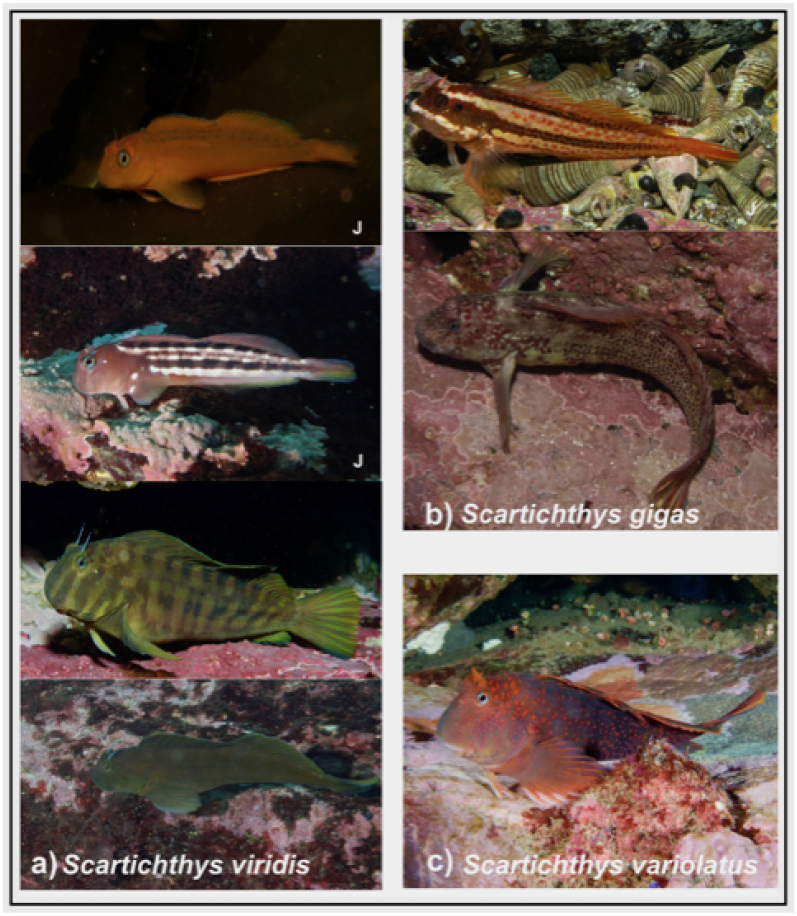
In situ photographs of juveniles (J) and adults of a) Scartichthys viridis; b) S. gigas and c) S. variolatus. All photographs taken by A. Perez-Matus.

## 5 CONCLUSION

This study presents evidence that S. crapulatus should be synonymized with *S. viridis*, resolving a 30-year-old discord among taxonomists. We also show that specimens displaying the colouration pattern described in 2016 (Mndez-Abarca & Mundaca, 2016) are juveniles of *S. viridis* which use different habitats (mainly occupying subtidal kelp sporophytes) than do adults. This study is a novel example of how molecular genetics can help to address the problem of species delimitation. We contribute to the clarification of the systematics of the Scartichthys genus, reconciling morphological, distributional, colouration and molecular evidence. We also show that the Juan Fernandez endemic species is relatively young and likely to be a neoendemic species.

## ACKNOWLEDGEMENTS

We thank Alex Gamarra, Sebastin Bez, Rodrigo Muoz and Italo Fernndez for their help in sampling specimens, Catalina Ruz, Ignacio Valverde and Sarai Morales for their help in processing samples in the laboratory. We thank Emily C. Giles for her technical assistance in the molecular laboratory and help improving the English of the manuscript. Finally, we thank N. K. Lujan and two anonymous reviewers for providing constructive reviews of earlier versions of the manuscript.

## AUTHOR CONTRIBUTIONS

EDT, HHS, PSA, MFL, and APM conceived the study; PSA, MFL, and APM acquired the funding; HHS, MFL, and APM collected the field data; EDT, HHS, PSA, and APM produced the data; EDT, HHS, PSA, and APM analysed the data; and all authors contributed to the writing and approved the final version of the manuscript.

## Funding Information

This research was funded by FONDECYT Grant # 1151094, FONDECYT Grant # 1210216 Proyecto Insercin Acadmica (PIA, 2731-010-81 of the P. Universidad Catlica de Chile to APM.) Collections of juvenile specimens in alcohol were funded by FONDECYT Grant # 1150296 grant to MFL.

## Notes

### Competing Interest Statement

The authors have declared no competing interest.

### Summary of Updates

Second round of revisions - improving English

