## Supplementary for "DNA reconciles morphology and colouration in the drunk blenny genus *Scartichthys* (Teleostei: Blenniidae) and provides insights into their evolutionary history"

Erwan Delrieu-Trottin^1,2^ | Hans Hartmann Salvo^3^ | Pablo Saenz Agudelo^4^ | Mauricio F. Landaeta^5,6^ | Alejandro Pérez Matus^3^

### **TABLE S1** Sampling locality, total length (*T_L_*), number of dorsal fin rays, number of dentary incisors (DI), coloration pattern in life (color in alcohol to thirteen specimens) and species name according to Williams (1990) and Méndez-Abarca & Mundaca, (2016) for each specimen, together with respective GenBank accession numbers for COI.

| **Specimen** | **Locality** | ***T_L_* (mm)** | **No. rays** | **No. incisors** | **Coloration pattern** | **Current species name** |
| --- | --- | --- | --- | --- | --- | --- |
| **P7** | S.J.M. | 97 | - | - | Orange-brown dots on posterior half of body | *S. crapulatus* |
| **P10** | S.J.M. | 97 | - | - | Dark-light bluish green | *S. viridis* |
| **P9** | S.J.M. | 98 | - | - | Reticulated bar-stained | *S. gigas* |
| **P6** | S.J.M. | 101 | - | - | Reticulated bar-stained | *S. gigas* |
| **P4** | S.J.M. | 115 | - | - | Uniform orange-brown | *S. gigas* |
| **P5** | S.J.M. | 124 | - | - | Dark-light bluish green | *S. viridis* |
| **P3** | S.J.M. | 128 | - | - | Reticulated | *S. gigas* |
| **P8** | S.J.M. | 141 | - | - | Reticulated | *S. gigas* |
| **P2** | S.J.M. | 143 | - | - | Reticulated | *S. gigas* |
| **P1** | S.J.M. | 156 | - | - | Reticulated | *S. gigas* |
| **A8** | Antofagasta | 52 | 18 | 68 | Uniform orange-brown | *S. gigas* |
| **A9** | Antofagasta | 56 | 17 | 58 | Uniform orange-brown | *S. gigas* |
| **A7** | Antofagasta | 66 | 17 | 83 | Uniform orange-brown | *S. gigas* |
| **A5** | Antofagasta | 70 | 17 | 103 | Uniform orange-brown | *S. gigas* |
| **A3** | Antofagasta | 93 | 18 | 58 | Two bar front head covered | *S. gigas* |
| **A6** | Antofagasta | 94 | 17 | 52 | Two bar front head covered | *S. gigas* |
| **A4** | Antofagasta | 127 | 17 | 64 | Reticulated | *S. gigas* |
| **A2** | Antofagasta | 175 | 18 | 113 | Dark-light bluish green | *S. viridis* |
| **A1** | Antofagasta | 246 | 17 | 60 | Reticulated | *S. gigas* |
| **M6** | Montemar | 49 | - | - | Whitish brown (color in alcohol) | - |
| **M3** | Montemar | 51 | - | - | Whitish brown (color in alcohol) | - |
| **M1** | Montemar | 51 | - | - | Whitish brown (color in alcohol) | - |
| **M2** | Montemar | 51 | - | - | Whitish brown (color in alcohol) | - |
| **M7** | Montemar | 52 | - | - | Whitish brown (color in alcohol) | - |
| **M4** | Montemar | 54 | - | - | Whitish brown (color in alcohol) | - |
| **M5** | Montemar | 55 | - | - | Whitish brown (color in alcohol) | - |
| **Q3** | Quintay | 50 | 18 | 93 | Uniform orange-brown | *S. gigas* |
| **Q2** | Quintay | 92 | 18 | 97 | Uniform orange-brown | *S. gigas* |
| **Q1** | Quintay | 103 | 18 | 109 | Uniform orange-brown | *S. gigas* |
| **EQ3** | El Quisco | 51 | - | - | Whitish brown (color in alcohol) | - |
| **EQ1** | El Quisco | 52 | - | - | Whitish brown (color in alcohol) | - |
| **EQ4** | El Quisco | 54 | - | - | Whitish brown (color in alcohol) | - |
| **EQ2** | El Quisco | 59 | - | - | Whitish brown (color in alcohol) | - |
| **PT5** | Punta de Tralca | 58 | 18 | 91 | Orange-brown dots on posterior half of body | *S. crapulatus* |
| **PT6** | Punta de Tralca | 94 | 17 | 108 | Orange-brown dots on posterior half of body | *S. crapulatus* |
| **PT3** | Punta de Tralca | 101 | 18 | 107 | Uniform orange-brown | *S. gigas* |
| **PT2** | Punta de Tralca | 104 | 18 | 118 | Uniform orange-brown | *S. gigas* |
| **PT4** | Punta de Tralca | 108 | 18 | 115 | Orange-brown dots on posterior half of body | *S. crapulatus* |
| **PT1** | Punta de Tralca | 214 | 18 | 119 | Dark-light bluish green | *S. viridis* |
| **LC16** | Las Cruces | 91 | 18 | 105 | Orange-brown dots on posterior half of body | *S. crapulatus* |
| **LC14** | Las Cruces | 92 | 19 | 104 | Orange-brown dots on posterior half of body | *S. crapulatus* |
| **LC3** | Las Cruces | 94 | 18 | 108 | Orange-brown dots on posterior half of body | *S. crapulatus* |
| **LC18** | Las Cruces | 95 | 18 | 112 | Orange-brown dots on posterior half of body | *S. crapulatus* |
| **LC17** | Las Cruces | 96 | 18 | 84 | Orange-brown dots on posterior half of body | *S. crapulatus* |
| **LC1** | Las Cruces | 98 | 18 | 113 | Orange-brown dots on posterior half of body | *S. crapulatus* |
| **LC10** | Las Cruces | 101 | 18 | 110 | Orange-brown dots on posterior half of body | *S. crapulatus* |
| **LC2** | Las Cruces | 101 | 18 | 113 | Orange-brown dots on posterior half of body | *S. crapulatus* |
| **LC15** | Las Cruces | 103 | 18 | 109 | Orange-brown dots on posterior half of body | *S. crapulatus* |
| **LC5** | Las Cruces | 105 | 18 | 116 | Orange-brown dots on posterior half of body | *S. crapulatus* |
| **LC9** | Las Cruces | 107 | 18 | 113 | Orange-brown dots on posterior half of body | *S. crapulatus* |
| **LC8** | Las Cruces | 107 | 18 | 115 | Orange-brown dots on posterior half of body | *S. crapulatus* |
| **LC11** | Las Cruces | 110 | 18 | 111 | Orange-brown dots on posterior half of body | *S. crapulatus* |
| **LC4** | Las Cruces | 112 | 18 | 115 | Orange-brown dots on posterior half of body | *S. crapulatus* |
| **LC6** | Las Cruces | 132 | 18 | 112 | Orange-brown dots on posterior half of body | *S. crapulatus* |
| **LC7** | Las Cruces | 134 | 18 | 123 | Orange-brown dots on posterior half of body | *S. crapulatus* |
| **LC13** | Las Cruces | 143 | 18 | 113 | Orange-brown dots on posterior half of body | *S. crapulatus* |
| **LC12** | Las Cruces | 155 | 18 | 116 | Orange-brown dots on posterior half of body | *S. crapulatus* |
| **IRC8** | Isla R. Crusoe | 77 | - | - | Circular pale spots in head and body (color in alcohol) | *S. variolatus* |
| **IRC9** | Isla R. Crusoe | 82 | - | - | Circular pale spots in head and body (color in alcohol) | *S. variolatus* |
| **IRC7** | Isla R. Crusoe | 105 | 18 | 84 | Circular red spots in head and body | *S. variolatus* |
| **IRC5** | Isla R. Crusoe | 124 | 18 | 93 | Circular red spots in head and body | *S. variolatus* |
| **IRC6** | Isla R. Crusoe | 137 | 18 | 89 | Circular red spots in head and body | *S. variolatus* |
| **IRC2** | Isla R. Crusoe | 157 | 18 | 94 | Circular red spots in head and body | *S. variolatus* |
| **IRC3** | Isla R. Crusoe | 157 | 18 | 95 | Circular red spots in head and body | *S. variolatus* |
| **IRC1** | Isla R. Crusoe | 188 | 18 | 95 | Circular red spots in head and body | *S. variolatus* |
| **IRC4** | Isla R. Crusoe | 206 | 19 | 93 | Circular red spots in head and body | *S. variolatus* |

### **TABLE S2** List of species and GenBank accession numbers (mainly from Hundt & Simons, 2018)for molecular loci used to construct the time‐calibrated phylogeny of *Scartichthys* spp.

| **Species** | **COI** | **ENC1** | **myh6** | **ptr** | **sreb2** | **tbr1** |
| --- | --- | --- | --- | --- | --- | --- |
| *Scartichthys gigas* | *forthcoming* | MG779111 | MG779150 | MG779181 | MG779032 | MG778946 |
| *Scartichthys variolatus* | *forthcoming* | MG7791112 | MG779151 | MG779182 | MG779033 | MG778947 |
| *Scartichthys viridis* | *forthcoming* | KF678531 | - | KF678719 | MG779030 | KF678816 |
| *Cirripectes quagga* | MH707862 | KF678520 | KF678592 | KF678683 | MG779034 | KF678777 |
| *Exallias brevis* | MK657660 | KF678520 | KF678617 | KF678709 | MG779029 | KF678806 |
| *Ophioblennius macclurei* | HQ168577 | KF678527 | KF678624 | KF678716 | MG779017 | KF678813 |

### **FIHURE S1** Magnified images 115 % for specimen PT1 (214 mm TL) of dark-light bluish green coloration (a) and zoomed at 82 % for specimen Q1 (103 mm TL) (b) showing the presence of circular orange spots (dots), difficult or impossible to see without a magnifying lens. Orange, brown to reddish dots (D) were observed for all specimens displaying the “dark-light bluish green” color pattern while only part of the specimens (A9, Q1, Q3 and PT2) displaying a “uniform orange brown” color pattern had such orange-reddish dots. Length of scale bar = 1 cm.

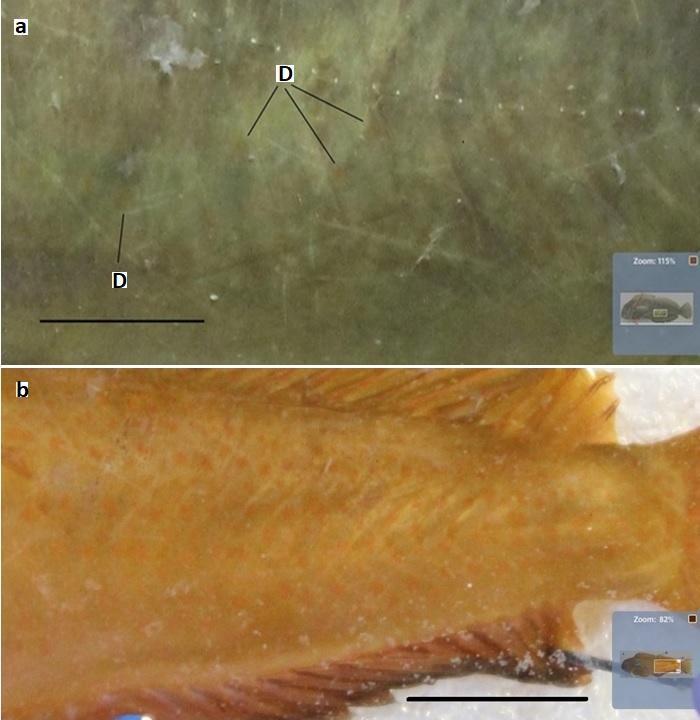
